# Comparison of Approaches to Transcriptomic Analysis in Multi-Sampled Tumors

**DOI:** 10.1101/2021.05.11.443668

**Authors:** Anson T. Ku, Scott Wilkinson, Adam G. Sowalsky

## Abstract

Intratumoral heterogeneity is a well-documented feature of human cancers associated with outcome and treatment resistance. However, a heterogeneous tumor transcriptome contributes an unknown level of variability to analyses of differentially expressed genes that may contribute to phenotypes of interest, including treatment response. Although current clinical practice and the vast majority of research studies use a single sample from each patient, decreasing costs in sequencing technologies and computing costs have made repeated-measures analyses increasingly economical. Repeatedly sampling the same tumor increases the statistical power of differentially expressed gene analysis that is indispensable towards downstream analysis and also increases ones understanding of within-tumor variance that may affect conclusions. Here, we compared five different methods for analyzing gene expression profiles derived from repeated sampling of human prostate tumors in two separate cohorts of patients. We also benchmarked the sensitivity of generalized linear models to linear mixed models for identifying differentially expressed genes contributing to relevant prostate cancer pathways based on a ground truth model.

## INTRODUCTION

Large-scale pan-cancer genomic analyses such as the Pan-Cancer Atlas [1] and the Pan-Cancer Analysis of Whole Genomes [2] have firmly established the heterogeneous and diverse nature of many primary tumors. Indeed, deep sequencing experiments have revealed that most cancers harbor complex phylogenies with distinct driver and passenger alterations, such that one region of a cancer may demonstrate a different phenotype than another part of the same tumor [3]. This spatial tumoral heterogeneity not only creates challenges for clinical diagnostics but also adversely impacts clinical outcome [4]. While the clinical convention of biopsying tumors to confirm diagnoses has historically been based on a single sample of tissue for most cancers, the regional histologic heterogeneity of prostate cancer lends itself to multiple sampling for a final diagnosis [5-8]. Paradoxically, prostate cancer molecular diagnostic tests, like many others, still rely on a single biopsy sample; the delicate balance between expensive and comprehensive testing is justified by treating the most aggressive-appearing tumor when given a choice [9]. Given the extent of tumor heterogeneity, however, sampling for research purposes has recapitulated diverse genomic and genotypic results across multiple samples of the same human tumor, which can similarly be reflected in discordant within-patient diagnostic scores [5-8].

For studies of the tumor transcriptome, measures of differentially-expressed genes (DEGs) are used to discover relationships between biological activity and phenotypes of interest, such as response to drug treatment or the impact of genomic alterations. With multiple-sampling strategies increasingly common due to lower costs of sequencing and computational power, key concerns linger pertaining to the consolidation or aggregation of multiple sources of (valuable) information from the same biological sample (*i*.*e*. a patient) with an outcome that is ascribed to the whole patient, and thus all samples as well [10-13]. For example, how much variance is acceptable to lose in the process of reconciling an evolutionarily and phenotypically diverse tumor to a discrete outcome? Perhaps, and more importantly, what measures related to outcome harbor intrinsically more variance deemed by repeated measures of the tumor?

For RNA sequencing, there is an underlying assumption that biological variability is often much greater between samples of different individuals than from within the same individual. Numerous bioinformatic solutions have been developed to assess the relationship between gene expression and outcome, often within the context of well-defined biological model systems with negligible within-individual variance and highly reproducible analytical measures such as RNA-seq. The most common tools include EdgeR, Limma/Voom, and DESeq2, and these tools are frequently used to compare phenotypes with a defined number of biological replicates for each phenotype, such that each replicate is given equal weighting using generalized linear models [14-16]. If applied to multiple samples of the same individual, options thus would include aggregating all within-individual data to individual-level values, or weighting each sample equally which would skew observations. An approach using DESeq2 would collapse pseudo-replicates by summing RNA-seq counts, which reduces the information available to discern fixed effects (variables that are constant across individuals, like treatment outcome) from random effects (categorical variables like somatic mutation status). Although Limma/Voom can accommodate the modeling of a single random effect, this would assume there are no other sources of variance to model, which is rarely the case when working with human biospecimens, especially if batch effects are considered. To wit, when extended to translational cancer research, modeling the transcriptomes of patient samples that are acquired or compared based innumerable salient phenotypes requires the inclusion of both fixed and random effects, which can be accommodated in linear mixed models. The R package for linear mixed models, lme4 [17] is used by the Dream package [18] for repeated-sampling analysis. However, its performance with real-world tumor data benchmarked against conventional analyses is unknown.

Here, we consider three potential genotypes based on somatic mutations that are relevant to prostate cancer (*TMPRSS2-ERG* fusion, *PTEN* loss, and 18q deletion) and the sampling of tumors from patients with those genotypes. We explore the impact of five different analytical methods on the differential expression analysis from two prostate cancer cohorts in which repeated-sampling design is a consideration, in which four of the methods collapse the specimens from the same lesion into a single sample (collectively termed aggregation) while the fifth method employs a linear mixed model. By comparing between-sample individuals of each cohort with known heterogeneous cancer somatic phenotypes to “gold standard” prostate cancer transcriptomes from The Cancer Genome Atlas (TCGA), we show that linear mixed models demonstrate superior sensitivity for identifying differentially expressed genes when controlling for strong heterogeneous confounders such as *PTEN* loss. Aggregation approaches show similar sensitivity to linear mixed models when working with fewer samples per individual or when working with phenotypes that result in little within-patient variability. As such, comparing the sensitivity of these methods further reveals sources of variance depending on target phenotypes and is a practical exercise for identifying occult causes of heterogeneity within a cohort. We thus provide this computational code as a resource for translational cancer researchers exploring heterogeneity within their own datasets.

## MATERIALS AND METHODS

### Patient cohorts

Cohort 1 consists of tumor samples from 37 patients diagnosed with intermediate to high-risk prostate cancer and subsequently treated with neoadjuvant hormone therapy and radical prostatectomy [8]. These tumor samples were acquired from MRI-targeted biopsies prior to the onset of therapy and guided by immunohistochemistry prior to laser capture microdissection. The manuscript by Wilkinson et al. [8] describes this cohort and each laser capture microdissected tumor focus.

Cohort 2 consists of tumor samples from 39 patients diagnosed with intermediate to high-risk prostate cancer and subsequently treated by radical prostatectomy [19]. These tumor samples were acquired from tissue punches of radical prostatectomy tissue, guided by a pathologist’s determination of low grade or high grade cancer regions. The manuscript by Ciampi et al. [19] describes this cohort and each tumor region.

### Raw sequence data processing

Paired-end RNA-seq FASTQ files for 137 tumor samples (cohort 1), 66 tumor samples (cohort 2), and 494 tumor cases (TCGA-PRAD) were downloaded from dbGaP (cohort 1), SRA (cohort 2), and the NCI Genomic Data Commons (TCGA-PRAD) and trimmed using Trimmomatic [20] version 0.36 using Illumina adaptor sequences with the parameter 3:50:10 MINLEN:36. Gene-level counts were estimated from paired-end reads aligned to version hg19 of the human genome using RSEM [21] version 1.3.2 as a wrapper around STAR [22] version 2.7.0f. Candidate gene fusions were detected using deFuse [23] version 0.8.1.

Paired-end BAM files corresponding to exome capture libraries for the matched samples in cohort 1 and cohort 2 were downloaded and converted to FASTQ files using GATK SamToFastq. FASTQ files were then re-aligned with the Burrows Wheeler Aligner [24] BWA-MEM version to version hg19 of the human genome (b37 with decoy chromosomes). The SAM alignment files were coordinate-sorted and duplicate-marked using PICARD version 2.18.27 SortSam and MarkDuplicates then quality score recalibrated using version 4.1.3.0 of the Genome Analysis Toolkit (GATK) BaseRecalibrator and ApplyBQSR. Mutation calling for ascertaining *PTEN* status was performed using MuTect2 (part of the GATK4 package), first by running MuTect2 in tumor-only mode on all of the normal tissue BAM files individually, with the parameters disable-read-filter set to MateOnSameContigOrNoMappedMateReadFilter and max-mnp-distance set to 0. The “padded” bait BED file (Agilent Human All Exon V7 for Cohort 1 and Agilent Human All Exon V5+UTR for Cohort 2), was used as the interval file of covered regions for all MuTect2 and filtering steps. Each of the output VCF files from this analysis of normal BAMs was merged into a database using GATK GenomicsDBImport with the setting merge-input-intervals set to true, and then generating a panel of normal from the database using GATK CreateSomaticPanelOfNormals. MuTect2 was run in somatic mode on each tumor-normal BAM pair and the panel of normals with af-of-alleles-not-in-resource set to 0.0000025 to exclude sites present in gnomAD and disable-read-filter set to MateOnSameContigOrNoMappedMateReadFilter. GetPileupSummaries and CalculateContamination were used on each tumor BAM file and the resultant contamination table was used to filter somatic mutations using FilterMutectCalls. CollectSequencingArtifactMetrics and FilterByOrientationBias were used to further filter mutations for 8-oxoG artifacts using the settings - AM G/T -AM C/T. These pass-filter mutations were then functionally annotated using Oncotator version 1.9.70 (database version April052016).

*PTEN* copy number changes were called following whole-genome assessment of somatic copy number alterations (SCNAs) across genomic intervals specified by the Agilent library design BED file with variable resolution depending on bait spacing. The design BED file was preprocessed with GATK PreprocessIntervals, with bin-length set to 0 and interval-merging-rule set to OVERLAPPING_ONLY. The BED file was also annotated with GC content using AnnotateIntervals. These interval files were used in all copy number calling steps. Read counts were first obtained from all tumor and normal BAM files using GATK CollectReadCounts. The normal read count files were compiled into a panel of normals using GATK CreateReadCountPanelOfNormals, excluding cases that had read depth coverage outside the 95% confidence interval for the dataset. The panel of normals was used to smooth read counts across all samples using GATK DenoiseReadCounts. Normal BAM files were then processed with GATK CollectAllelicCounts to identify regions of potential LOH. GATK CollectAllelicCounts was also applied to each tumor BAM file. GATK ModelSegments used smoothed read counts from each tumor BAM along with the paired normal/tumor allelic counts for generating copy number estimates.

Calls of *PTEN* somatic copy number changes were performed using GISTIC 2.0.23 on the GenePattern platform (module version 6.15.28). The SEG file from GATK ModelSegments was used as input with the following parameters: focal length cutoff set to 0.50, gene GISTIC set to yes, confidence level set to 0.90, cap value set to infinite, broad analysis set to on (to call arm-level events), max sample segments set to 10,000, arm peel set to no, and gene collapse method set to extreme. Post-GISTIC log_2_-copy number ratio threshold values were used for calling discrete gene-level calls as follows: > 1.3 = 2-copy gain; 0.1 to 1.3 = 1-copy gain; -1.3 to -0.1 = 1-copy loss; < -1.3 = 2-copy loss. Chromosome 18q changes followed the same threshold scheme per tumor focus, using the convention that at least 50% of the chromosome arm (noncontiguously) must be affected.

### Genotype/phenotype determination

For all TCGA cases, the reported status of *TMPRSS2-ERG, PTEN*, and 18q were used as deposited on cBioPortal for the PanCancer Atlas analysis (comprising 489 tumors). Fusions involving *TMPRSS2* or *ERG* and other genes (such as *SLC45A3-ERG*) were excluded. Cases with undocumented 18q status, or gains to *PTEN* or 18q were also excluded. For cohort 1, *PTEN* status was determined jointly based on the status of anti-PTEN immunohistochemistry reported for each focus [8] and somatic copy number or mutation calls. For Cohort 2, *PTEN* status was determined based on somatic copy number or mutation calls. For cohort 1, *TMPRSS2-ERG* status was determined jointly based on the status of nuclear anti-ERG immunohistochemistry reported for each focus [8] and RNA-seq fusions determined by deFuse. For cohort 2, *TMPRSS2-ERG* status was determined based on RNA-seq fusions determined by deFuse supplemented with any observed interstitial deletions between *ERG* and *TMPRSS2* on chromosome 21 detected by somatic copy number analysis.

Aggregation of ground-truth phenotypes from foci to lesions and/or patients was as follows: a pre-determined threshold was set for a minimum number of foci to harbor that alteration to call a particular copy number event for the entire lesion. For lesions sampled by 7 or 8 foci, the threshold was 3; for 3, 4, 5 or 6 foci the threshold was 2, and for 1 or 2 foci the threshold was 1. For each focus, *PTEN* or 18q somatic copy number calls from GISTIC (*i*.*e*., -1, -2, 0, 1 or 2) were averaged across all foci from each lesion. If the absolute value of that average was greater than the pre-determined threshold, the averaged value was rounded to the nearest integer. For *PTEN* point mutations or the *TMPRSS2-ERG*, the lesion (cohort 1) or patient (cohort 2) was considered to harbor the mutation if any number of foci harbored it. Similarly, if any focus harbored PTEN loss or ERG expression by immunohistochemistry, the entire lesion (cohort 1) or patient (cohort 2) was considered to have harbored it. Lesions marked as the index lesion were used as patient-level data for cohort 1.

### Sample counts aggregation

Five (cohort 1) or four (cohort 2) different methods of transcriptomic analysis of multiple-sampled tumors were investigated as depicted in Figure 1. In the first method, concatenation, paired-end FASTQ files derived from samples of the same lesion or tumor were concatenated into a single pair of FASTQ files representing the entire lesion or tumor. These FASTQ pairs were then processed as above with RSEM and STAR, arriving at a single counts file per lesion or tumor. The remaining analyses were performed with sample counts obtained after alignment and count estimation as stated above. In the second method, averaging, counts from the same lesion were averaged across genes, creating a representative lesion-level transcriptome profile for each lesion or patient. In method three, RNA weighting, gene counts were normalized based on the weighted average of the specimen RNA input amounts into library preparation as seen in equation 1.

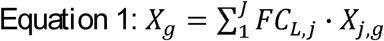

where *X*_*g*_ is the count of gene *g*, FC_j_ is defined as the fractional concentration of specimen *j* obtained from lesion *L* calculated as equation 2.

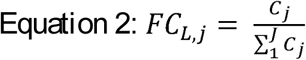

**Figure 1.**
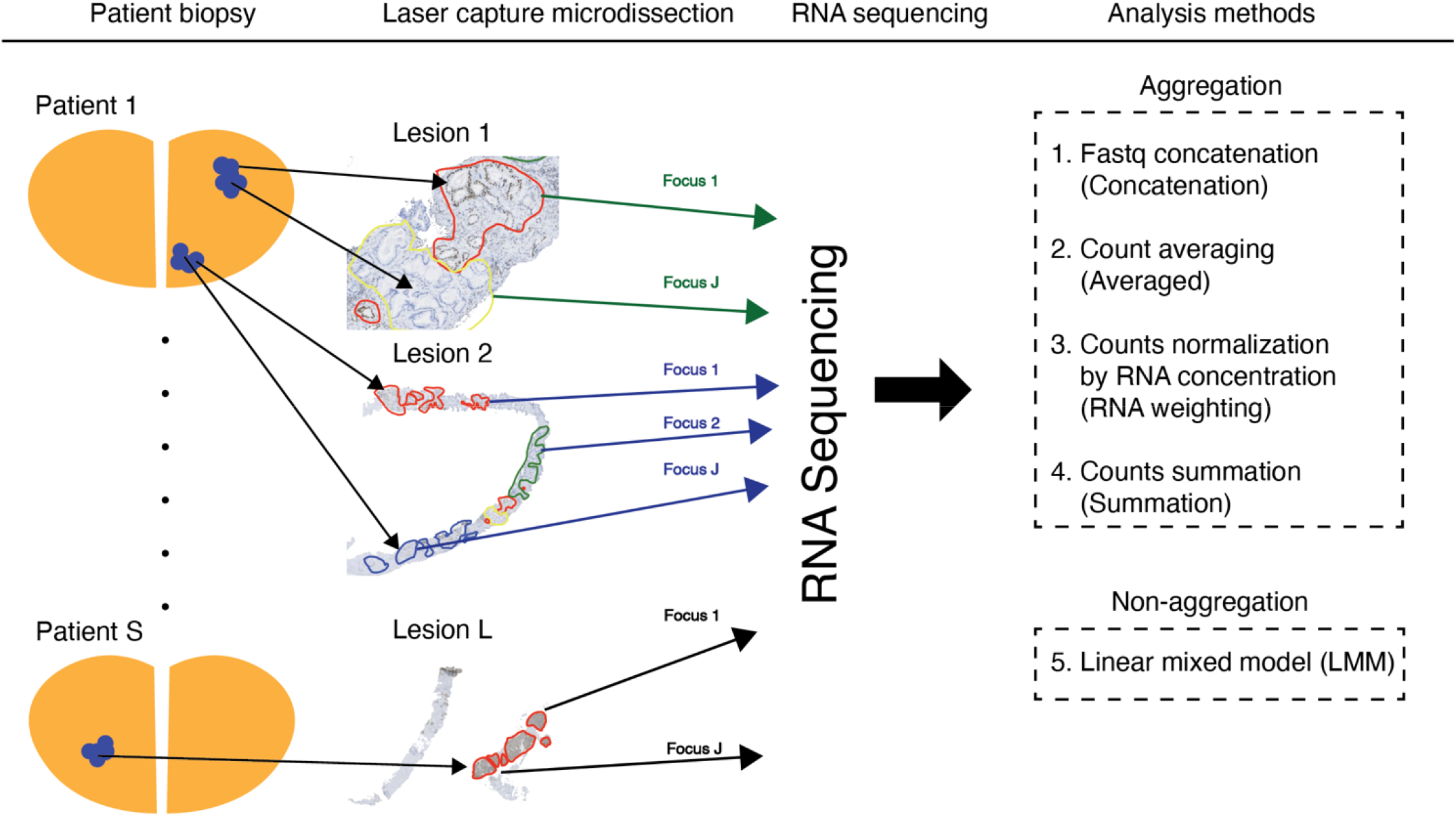
Schematic illustration of the work flow for cohort 1. Biopsies were obtained from one or more lesions per patient. For each lesion, one or more focus were laser micro-dissected (LCM) from biopsied materials. RNA sequencing were performed on each focus and the associated ERG and PTEN protein status were determined by immunohistochemistry of adjacent slides while chromosome 18q status were was from exome sequencing data. The gene expression data were analyzed by five different methods, FASTQ concatenation, counts averaged, RNA weighting, counts summation and linear mixed model.

Due to missing data, RNA weighting was only performed for cohort 1. In method four, Summation, counts of the same lesion/patient were added across genes to create a high coverage profile of the tumor’s transcriptome. Finally, in method five, the linear mixed model (LMM) was used with fixed and random effects to account for the inter-patient and inter-lesion variability.

### Differential gene expression analysis

Differential gene expression analysis for aggregated samples in cohorts 1 and 2 was performed using the DESeq2 package available in R [16]. Gene count matrices were prepared as described above using RSEM/STAR. Genes with zero counts in more than half of the samples were removed. Next, counts were normalized by the median ratio method from DESeq followed by differential expression analysis using either *TMPRSS2-ERG* fusion, *PTEN* loss or chr18q loss as fixed effect parameters. Genes with false discovery rate (*p*_adj_) less than 0.05 and log_2_ fold-change less than -1 or greater than 1 were considered differentially expressed.

### Linear mixed model analysis

Differential gene expression analyses using a linear mixed model were performed using the variancePartition package available in R [18]. First, gene counts were normalized by trimmed the mean method in edgeR. Next, the genomic alterations of interest were modeled as fixed effects while patients and their lesions were modeled as nested random effects in the LMM model using the formula: ∼ Fixed_effect + (1|Patient:Lesion) for cohort 1. For cohort 2, the genomic alterations of interest were modeled as fixed effects while patients were modeled as a random effect variable using the formula: ∼ Fixed_effect + (1|Patient).

### Derivation of *PTEN* loss, *TMPRSS2-ERG* and 18q-loss gene sets

Gene count matrices from TCGA-PRAD were annotated with *PTEN, TMPRSS2-ERG* and 18q status, and differential expression analysis based on each genotype was performed using DESeq2. After removing genes with zero counts in at least half of the samples, genes were normalized by the median of ratios method as described in DESeq2. Gene signatures were generated by performing differential expression analysis using DESeq2each genotype as an input parameter and accepting DEGs with *p*_adj_ < 0.05 by Wald test.

### Statistics

Similarities between groups of samples were determined using unsupervised hierarchical clustering (Euclidean distance). Null hypothesis tests of associations between affected and unaffected samples were performed using two-sided Fisher’s exact tests. Generalized linear models using the likelihood ratio test were used to compare gene expression between groups of singly-sampled tumors. Unless noted otherwise, an alpha of 0.05 was used as a significance threshold for all statistical analyses.

### Software

All mathematical analyses were performed using R version 4.0.2 and RStudio 1.3.

## RESULTS

### Transcriptomic consequences of genomic heterogeneity in prostate tumors

Despite its common genetic background, within-patient tumor heterogeneity manifests itself through variable histologies and genomic alterations, even by geographically nearby clusters of tumor cells. As a framework for assessing salient gene expression as a function of histologic or genomic heterogeneity, we selected three somatic alterations common to prostate cancer: genomic deletion or deleterious alterations to the tumor suppressor *PTEN*, loss of at least half of chromosome 18q, and the fusion of *TMPRSS2* and *ERG* on chromosome 21. Alterations to *PTEN* and 18q were determined by analysis of whole-exome sequencing data from matched samples (compared to benign controls), while *TMPRSS2-ERG* fusion was determined by the expression of nuclear ERG by immunohistochemistry.

We assessed the frequency of these alterations in two distinct cohorts. Cohort 1 was comprised of 137 laser capture microdissected (LCM) foci, sampled from 51 different biopsies targeting 50 distinct imaging-visible lesions from 39 patients, with 1-3 lesion per patient and 1-8 foci per patient [8]. Foci in cohort 1 were microdissected based on distinct histologic features, such as gland architecture, proximity to other tumor cells, and immunochemistry against common prostate cancer markers such as anti-ERG and anti-PTEN. Cohort 2 was comprised of 66 tumor punches from either high-grade or low-grade tumors from prostatectomy samples assessed without imaging, with 1-2 foci per patient [19]. Regions sampled in cohort 2 were determined solely on tumor aggressivity.

We first performed unsupervised clustering on gene expression data from each focus individually, in both cohorts, annotating each sample for *PTEN*, ERG, and 18q status. Of these three factors, clustering revealed that foci clustered most strongly based on ERG fusion status, and then by *PTEN* status, both in cohort 1 (Fisher’s exact test, adjusted *p* = 2.2×10^−23^; 1.5×10^−10^; and 1 respectively, *k* = 4; Fig. 2a) and in cohort 2 (Fisher’s exact test, adjusted *p* = 2.10^−5^; 0.041; and 1 respectively, *k* = 2; Fig. 2b). Although ERG fusion and *PTEN* loss showed significant per-focus co-occurrence in cohort 1 but not cohort 2 (Fisher’s exact test, *p* = 2.0×10^−9^ and 0.6, respectively), we thus assessed the inter- and intra-lesion variation of ERG, *PTEN* and chr18q alterations of cohort 1 and the intra-patient variation in cohort 2. *PTEN* loss had the highest inter- and intra-lesion variability of the three alterations in cohort 1 (Figs. 2A and 2B) and the highest intra-patient variability in cohort 2 (Figs. 2C and 2D), suggesting that the subclonal nature of major driver alterations such as *PTEN* loss could significantly affect tumor phenotypes with a patient.

**Figure 2.**
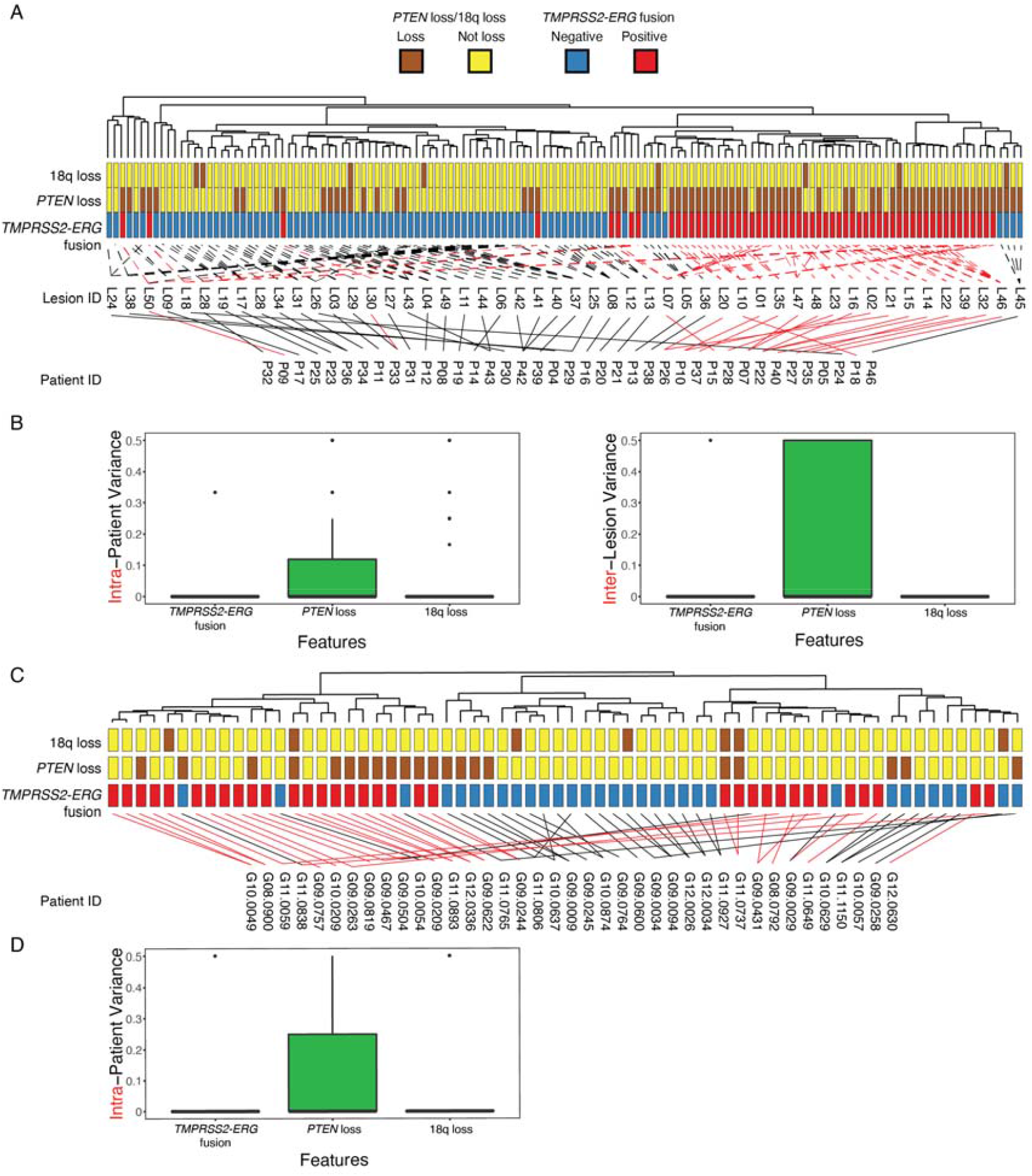
Summary of gene expression profile of cohort 1 and 2. **(A)** Unsupervised clustering of focus level gene expression data from cohort 1 with ERG, PTEN and chr18q loss annotation. The results revealed a strong tendency for focus to group based on ERG and PTEN status (Fisher’s exact test, adjusted *p* = 2.2×10^−23^ and 1.5×10^−10^, respectively) but not chr18q loss (*p* = 1). Corresponding patient identifier and lesion identifier were mapped below the dendrogram. Black lines corresponding to foci or lesions without ERG fusion were drawn connecting to lesions and patients while red lines corresponds foci or lesions with ERG fusion foci. **(B)** Assessment of ERG, PTEN and chr18q inter and intra lesion heterogeneity. PTEN loss remains the most variable feature within and between lesions in cohort 1. **(C)** Unsupervised clustering of focus level gene expression data from cohort 2 with ERG, PTEN and chr18q status annotation. The results revealed a strong tendency for focus to group based on ERG and PTEN status (Fisher’s exact test, adjusted *p* = 2.10^−5^ and 0.041, respectively) but not chr18q loss (*p* = G1). **(D)** Assessment of ERG, PTEN and chr18q intra lesion heterogeneity. PTEN loss remains the most variable feature within lesions in cohort 2.

### Transcriptomic consequences of somatic genomic alterations

Patient response to cancer therapies are often distilled to a binary value: remission or recurrence, response or failure, survival or mortality. Precision medicine workflows to predict such clinical responses most often rely upon a single sample from which various measures are taken. However, assessments of tumor heterogeneity (i.e. multiple region sampling) are exploratory in nature and for research purposes; given the geometric increase in cost for clinical assays and the challenges associated with interpreting multiple and potentially conflicting results from a single patient, repeated clinical assays of the same sample per patient are not feasible. Thus, the translatability of human oncology research depends upon a definitive outcome wherein the evidence supporting that result is a takes into account phenotypic variability due to heterogeneity.

We thus considered five different methods for deriving differentially expressed genes (DEGs) from our two cohort when working with tumors that were sampled multiple times: (1) averaging of counts within each case (averaged), (2) weighted averaging of counts based on RNA input amounts into library preparation (RNA weighting), (3) concatenation of FASTQ files prior to a single analysis per case (concatenation), (4) summation of counts (summation), and (5) use of a linear mixed model (LMM). DEGs were derived for each cohort and method based on the status of ERG (*TMPRSS2-ERG* fusion positive vs. negative), PTEN (reduced or lost vs. intact) and chromosome 18q (lost vs. intact). For aggregation methods 1-4, if a single sample was considered altered, the entire case was considered altered. For the LMM (method 5), *TMPRSS2-ERG, PTEN* and 18q status was specified as a fixed effect while cases were modeled as random effect.

Table 1 shows the number of statistically significant DEGs (*p*_adj_ < 0.05). Generally, the number of DEGs was dependent on the genomic alteration and the method of aggregation. *TMPRSS2-ERG* fusion, a categorically early event in the natural history of prostate tumors, yielded the largest number of DEGs, followed by *PTEN* loss (which tends to occur diffusely amongst major subclones) and chr18q loss (even more frequently subclonal), respectively. The number of significant DEGs derived from each method differed based on method and genomic feature analyzed. Amongst the individual DEGs, *ERG* ranked at the top three up-regulated genes by all five methods in cohort 1 and the top nine up-regulated genes in all four methods in cohort 2 (Table 2). Similarly, *PTEN* was found to be down-regulated in PTEN loss cases by all five methods in cohort 1 but not in cohort 2. For chr18q loss, DEGs associated with chr18q loss were not located in chr18q for both cohort 1 and 2. The complete lists of DEGs for each genotype are given in Supplementary Tables 1-3 (cohort 1) and Supplementary Tables 4-6 (cohort 2).

**Table 1.**
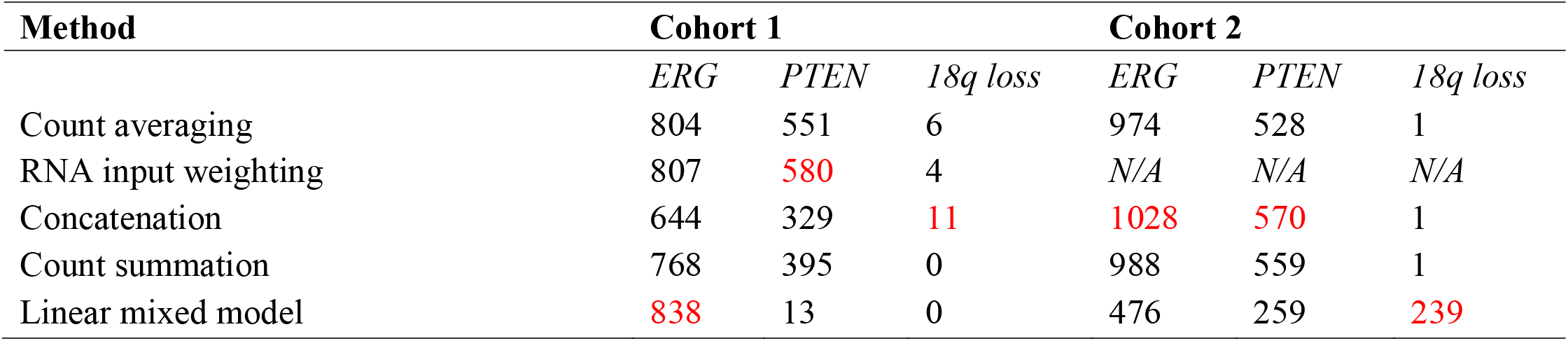
The total number of differentially expressed genes (DEGs) for each aggregation method per cohort, per genotype. The greatest number of DEGs for each genotype is shown in red.

**Table 2.**
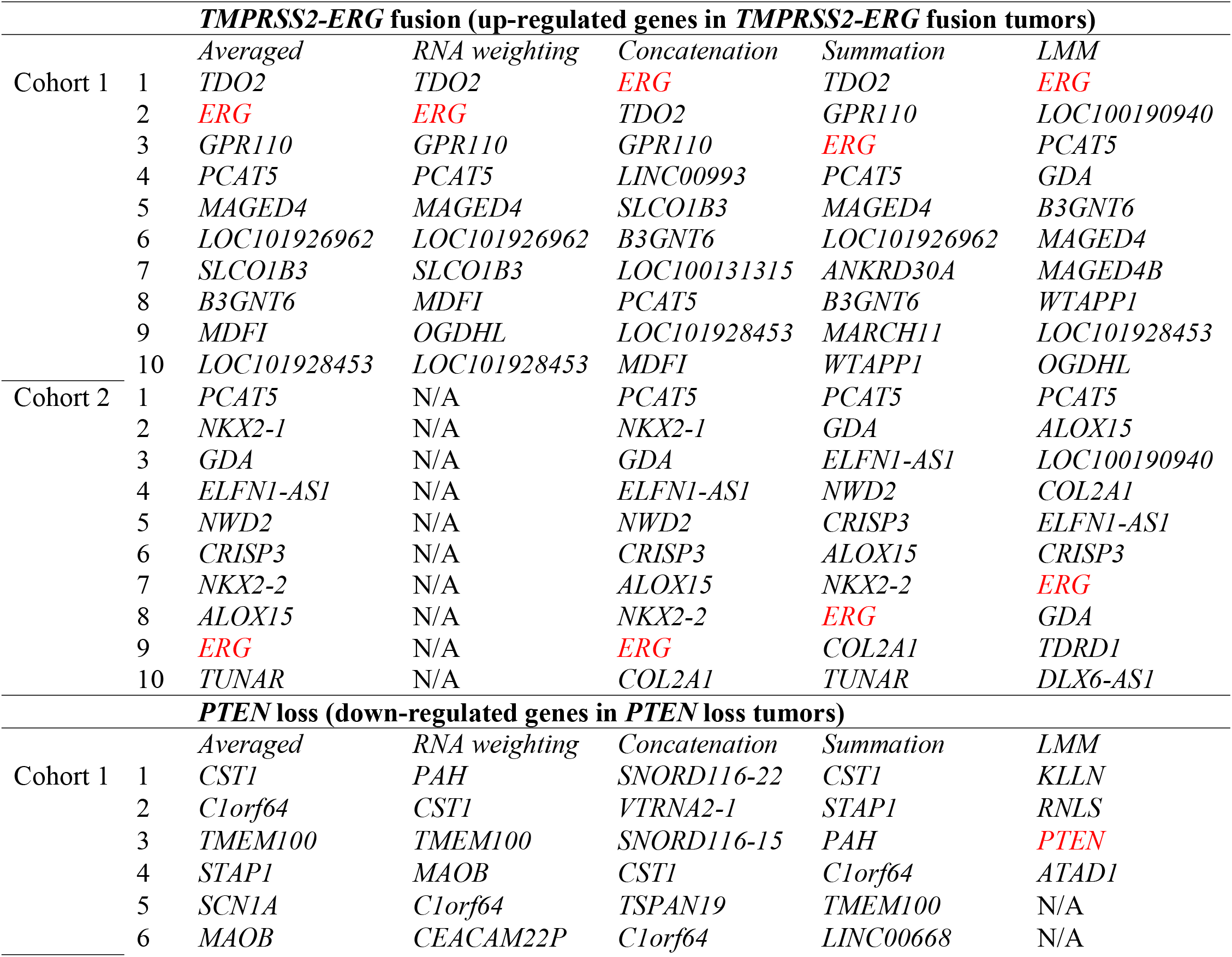
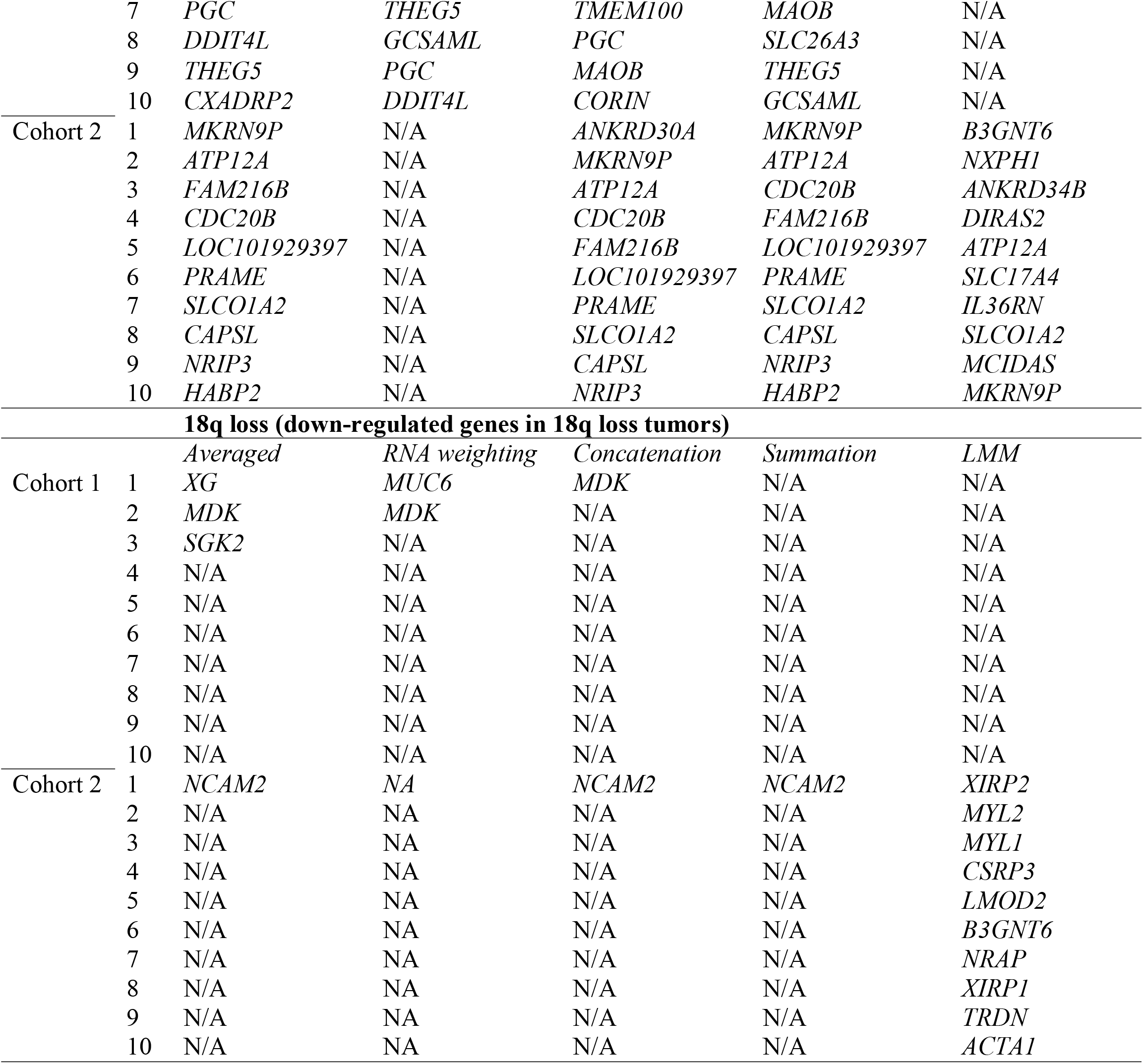
The top 10 up- or down-regulated genes for each aggregation method per cohort, per genotype, that pass the threshold of log_2_fold change of ±1 at *p*_adj_ < 0.05. *ERG*, which is expected to be an up-regulated gene in the *TMPRSS2-ERG* gene set, and *PTEN*, which is expected to be a down-regulated in the *PTEN* loss gene set, are shown in red for emphasis.

By set analysis, 393, 4 and 0 DEGs were shared between the five methods for *TMPRSS2-ERG* fusion, *PTEN* loss and chr18q loss, respectively in cohort 1 (Fig. 3A). Similarly, a total of 377, 143 and 0 genes were common for cohort 2 (Fig. 3B). To determine if the number of DEGs produced by different method was sensitive to alpha threshold, the number of unique DEGs obtained from all five methods were pooled and as a function of increasing alpha for all three genomic features in cohorts 1 (Fig. 4, top) and 2 (Fig. 4, bottom). We observed that by increasing alpha from 0.05 to 0.2, the number of detected DEGs did not increase appreciably in all three genotypes across both cohorts.

**Figure 3.**
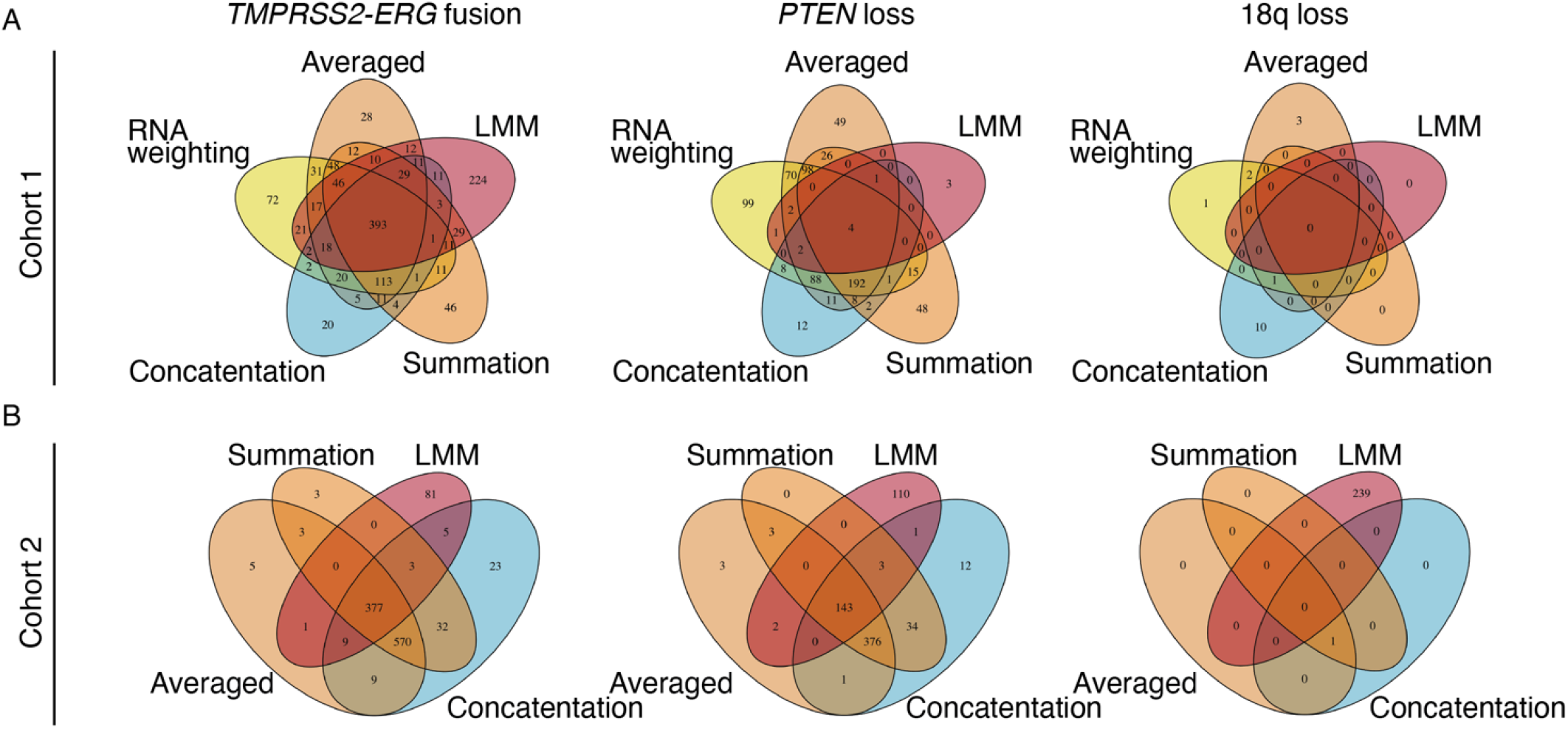
Set analysis of genes related to ERG fusion, PTEN loss and chr18q loss in cohort 1 and 2. **(A)** Venn diagram depicting the shared DEGs between each aggregation method in cohort 1. The number of DEGs common between all five methods are 393, 4 and 0 for ERG, PTEN loss and chr18q loss, respectively, in cohort 1. **(B)** Venn diagram depicting the shared DEGs between each aggregation method in cohort 1. The number of DEGs common between the five methods are 377, 143 and 0 for ERG, PTEN loss and chr18q loss, respectively, in cohort 2.

**Figure 4.**
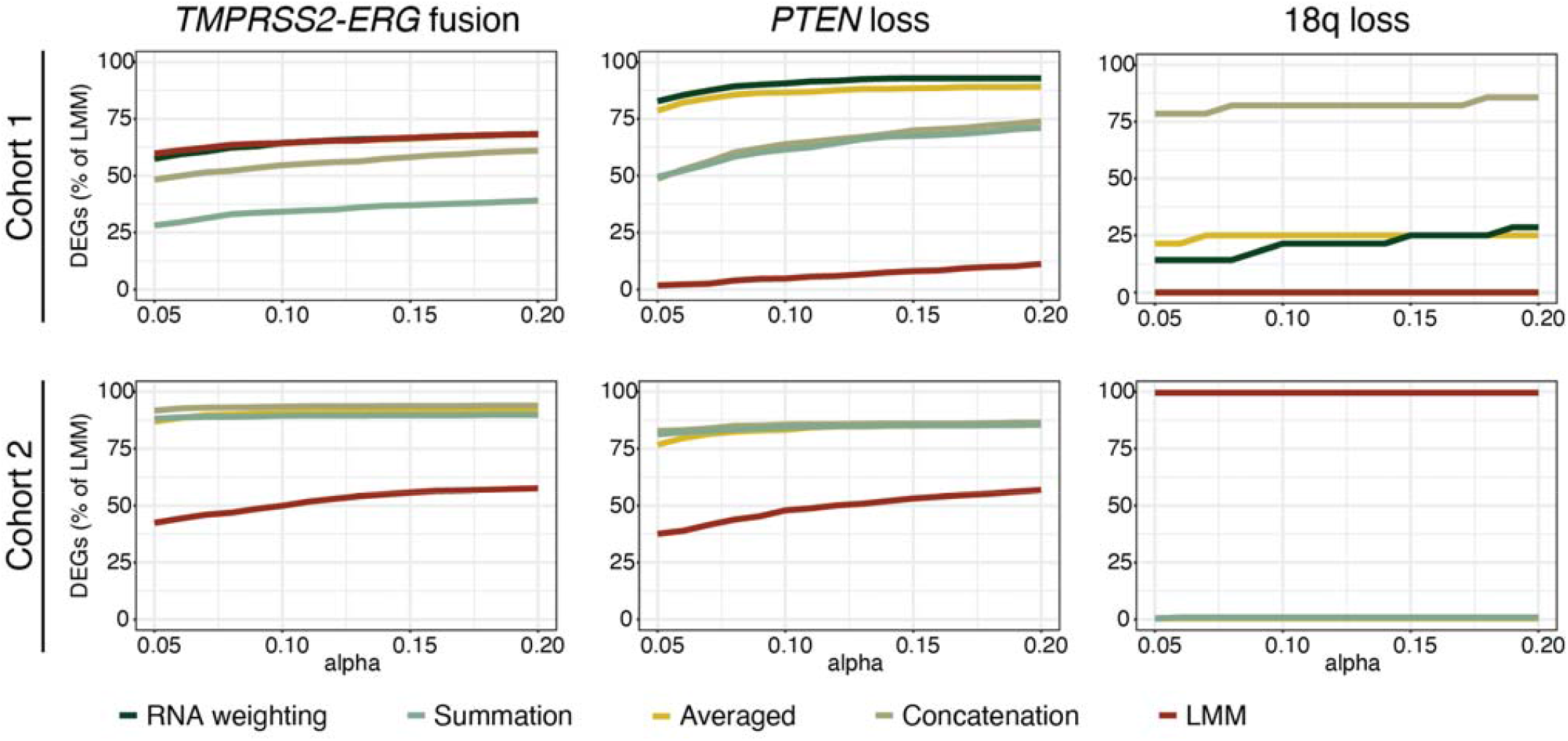
Sensitivity analysis of DEGs to alpha. For each genomic alteration analysis, the total number of DEGs from each method were pooled. The filtering threshold were set to 1 ≤ log_2_ fold-change ≥ 1 followed by increasing alpha to determine the percentage of significant DEGs. The increase in the alpha from 0.05 to 0.20 did not lead to a convergence in the number of DEGs across genotype in cohort 1. Sensitivity curves were grouped the closest in *TMPRSS2-ERG* fusion genotype but showed the largest magnitude difference in *PTEN* loss and chr18q loss. Averaged, concatenation and summation are consistently grouped together with increasing alpha from 0.05 to 0.20 in cohort 2. LMM did not converge with the other methods.

When we assessed the similarities between the five different methods using unsupervised clustering of the differential expression output, we noted that LMM clustered away from other methods for *PTEN* loss analysis of cohort 1 (Fig. 5A) and all genotypes in cohort 2 (Fig. 5B), suggesting that in cohort 1, the effect of *PTEN* loss on heterogeneous gene expression profiles within a patient was apparent due to the separately-captured samples. In particular, the clinical practice of determining an entire cancer being called PTEN-negative due to at least 5% of tumor cells showing PTEN reduction by IHC has a similar effect on heterogeneity when multiple samples from the same tumor are captured without microdissection.

**Figure 5.**
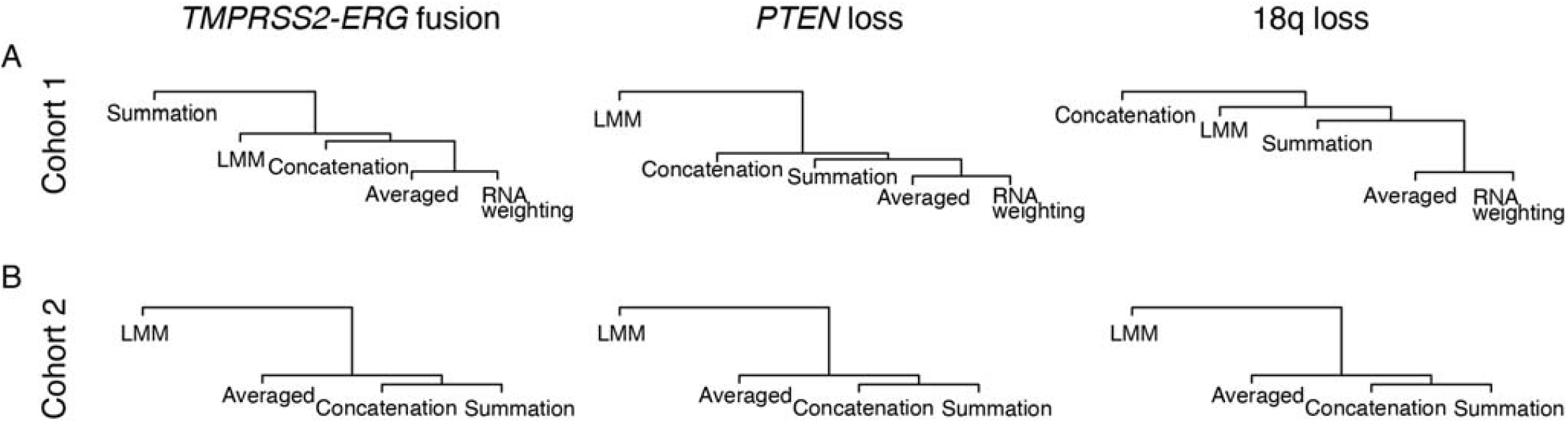
The relatedness of the five methods were assessed by hierarchical clustering of log_2_ fold-change from the five methods in cohort 1 (A) and four methods in cohort 2 (B). The type of genomic change is indicated above each tree.

At the sample level, *TMPRSS2-ERG* fusion and reductions in *PTEN* were commonly co-occurring in cohort 1, so we assessed the number of DEGs that could spill over from *TMPRSS2-ERG* into *PTEN* analysis in all five methods. The result revealed a total of 269, 283, 179, 188 and 4 genes that are common between *TMPRSS2-ERG* fusion and *PTEN* DEG analysis from counts averaging, RNA weighted averaging, concatenation of FASTQ files, counts summation and LMM methods respectively. To better understand the impact of within-patient variability and remaining variance after accounting for each genomic alteration on gene expression, we performed variance decomposition using results from LMM for both cohorts (Figure 6). Indeed, between-patient variance was a significant source of transcriptomic heterogeneity, explained by a median of 33-36% percent of variance in ERG fusion, PTEN loss and 18q loss gene expression in cohort 1 and 55-59% for cohort 2 respectively. In contrast, genomic alterations explained a median of 0% variance in both cohorts when accounted for as a fixed effect.

**Figure 6.**
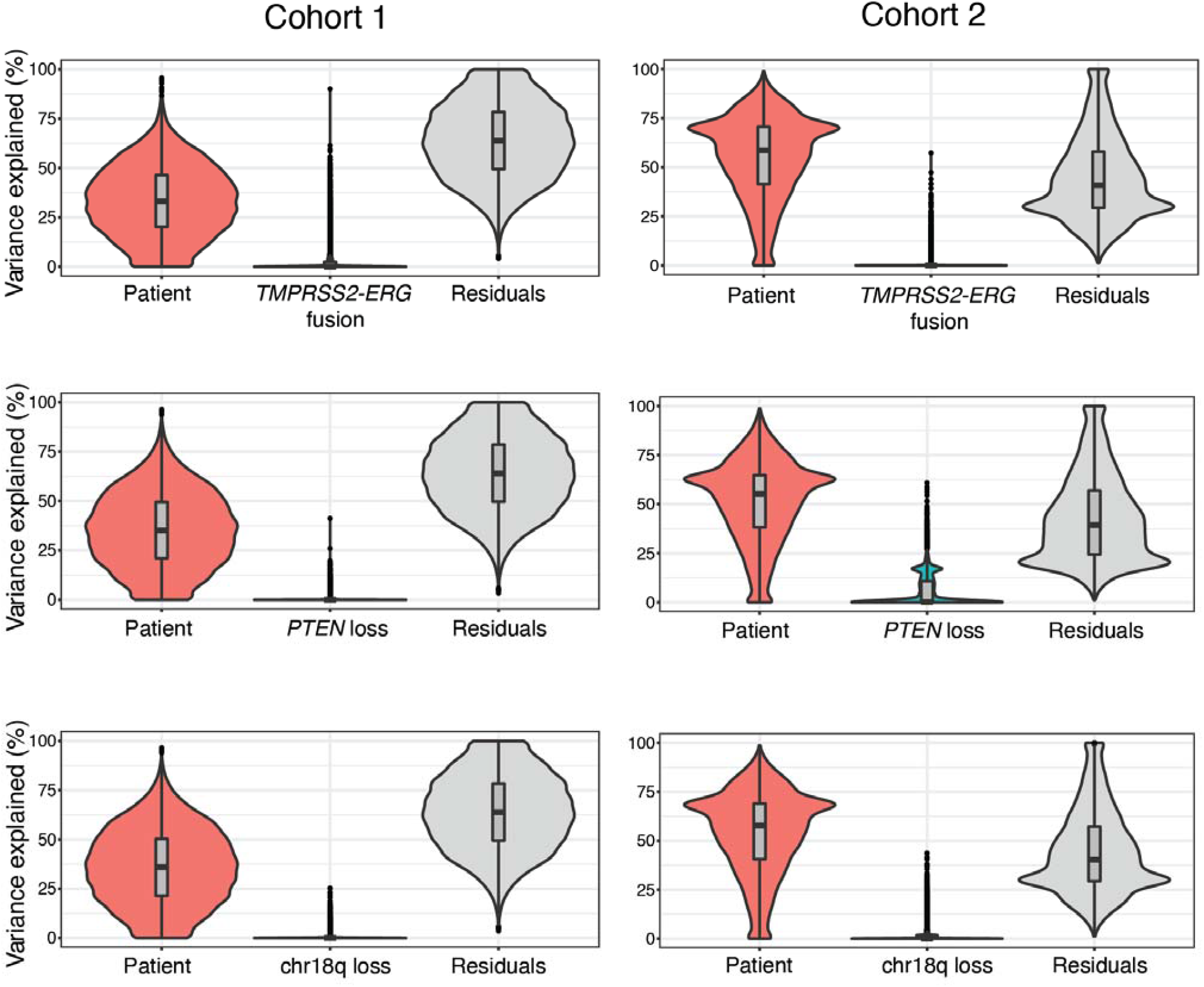
Decomposition of gene expression variation into patient, genotypic feature and residuals from cohort 1 (left) and cohort 2 (right). Patient, genomic features and residuals explain a median of 33%, 0%, and 64% for *TMPRSS2-ERG* fusion, 35%, 0% and 64% for *PTEN* loss and 36%, 0% and 64% for 18q loss respectively. For cohort 2, patient, genomic features and residuals can explain a median of 59%, 0% and 40% for ERG, 55%, 0% and 39% for PTEN loss and 58%, 0% and 40% for 18q loss.

### Comparison of differentially expressed genes to a standardized gene set

As blocking for each specific genomic alteration reduced variance in the LMM, we still have observed that multiple methods for deriving DEGs for each genotype produced overlapping but distinct results. Thus, a key question is whether one approach arrives at a more accurate representation of *TMPRSS2-ERG, PTEN*, or 18q-dependent genes. To benchmark these approaches, we next derived ground-truth gene sets based on *TMPRSS2-ERG* fusion, *PTEN* loss, and 18q loss by performing an integrated analysis of the prostate adenocarcinoma cancer genome atlas (TCGA-PRAD) cohort, where significantly up-regulated genes in the perturbed phenotype (*e*.*g. TMPRSS2-ERG* fusion-positive) comprised the gene set. Overall, *TMPRSS2-ERG* fusion calls were made in 478 cases (172 cases were positive for the fusion), *PTEN* status was determined in 427 cases (103 cases harbored *PTEN* loss-of-function mutations or copy number losses), and 18q status was called in 416 cases (with 61 cases harboring 18q losses). After determining the differentially expressed genes for each genotype, we applied a cutoff to keep the top 75 up-regulated genes for each gene set. The genes comprising these signatures are provided in Supplementary Table 7.

Finally, we performed gene set enrichment analysis (GSEA) to assess the sensitivity of each aggregation approach for cohorts 1 and 2 based on known alteration status to each corresponding gene set. Employing normalized enrichment score (NES) as a measure of sensitivity, the five methods produced a range of NES that was highly dependent on genomic alteration. As depicted in Figure 7, GSEA of ERG fusion produced the highest and statistically significant NES for both cohorts, ranging from 3.0 to 3.2 (cohort 1) and 3.0 to 3.4 (cohort 2). The methods that yielded the highest NES were concatenation for cohort 1 and summation for cohort 2 respectively. For *PTEN* loss, the NES were significant and ranged from 2.1 to 2.59 for cohort 1 and 2.88 to 3.12 for cohort 2. LMM performed the best for both cohorts. For chromosome 18q loss, LMM performed comparably to averaging in both cohorts, but did not approach the biologically significant threshold of ±2.

**Figure 7.**
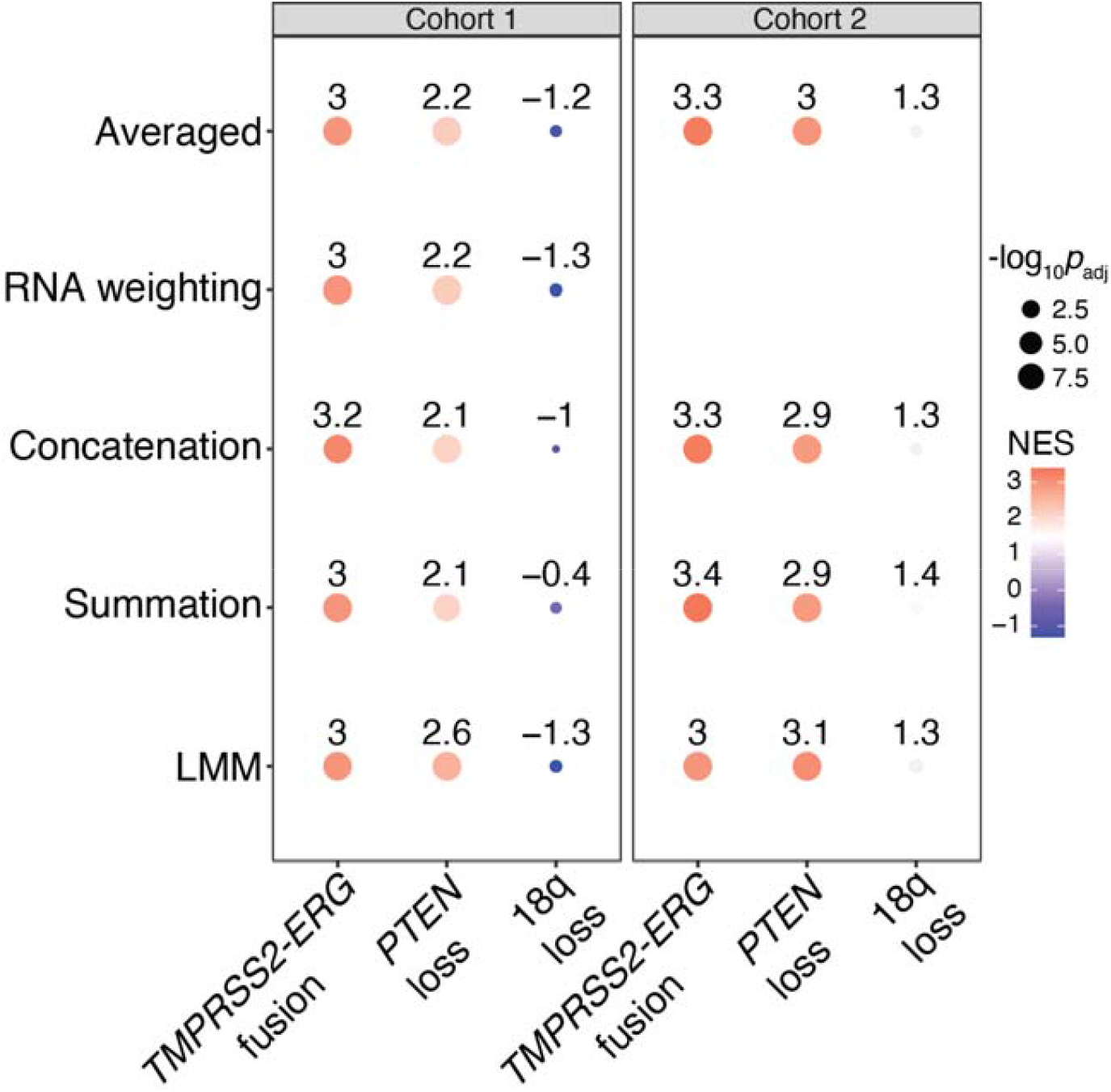
Bubble plot illustrating the normalized enrichment score (NES), colored from low (blue) to high (red) and -log_10_ adjusted *p* value (size of the dot) derived from gene set enrichment analysis of *TMPRSS2-ERG* fusion, *PTEN* loss and chr18q loss using differential expression analysis outputted from counts averaging, RNA weighting, concatenation, summation and linear mixed model (LMM).

## DISCUSSION

With the increasing recognition of tumor heterogeneity as a significant prognostic factor for patient outcome [3, 8, 25], experiments that account for tumor heterogeneity show promise to improve the statistical power of differential gene expression (DEG) analysis. Practically, the more frequently a tumor is measured, within-tumor variance for particular genes can indicate relative importance towards salient phenotypes and genotypes. Any intra- and inter-tumoral transcriptomic variance could be assessed using statistical models such as linear mixed model. However, the benefits to experimental designs such as these have yet to be fully explored, especially in the context of other aggregation methods that collapses multi-region samples into a single sample. Therefore, we investigated herein the impact of four methods for aggregation of RNA-seq counts and compared them to linear mixed modeling for detecting DEGs related to the important prostate cancer (PCa) genotypes of *TMPRSS2-ERG* fusion, *PTEN* loss and chr18q loss.

These three target genotypes range in frequency and relevance in prostate cancer, with *TMPRSS2-ERG* fusion generally being a very early event in PCa tumorigenesis, followed by *PTEN* loss and 18q loss [26, 27]. This is reflected in part by the clonal or subclonal nature of these events, and the potential ability to isolate pure populations of tumor cells using laser capture microdissection based on stains detecting these events. Due to microdissection in cohort 1, the specificity of transcriptome underpinning the phenotypic differences between *TMPRSS2-ERG* fusion-positive and negative tumors between cases was so great that all aggregation methods performed comparably (see Fig. 5A). In contrast, the loss of chr 18q which is likely a passenger event did not produce robust DEGs and GSEA normalized enrichment score (NES) from any methods. For *PTEN* loss in cohort 1, and all target genotypes in cohort 2, the DEGs derived by a linear mixed model (LMM) were substantially different than all other methods (see Fig. 5B). The simplest explanation for this observation is that a per-patient classification of a genotype such as “*PTEN* loss” demonstrates more heterogeneity across the tumor, as PTEN-intact and PTEN-reduced tumor cells harbored strikingly different inter-lesion and intra-lesion gene expression profiles. Additional evidence from variance decomposition supports the notion that majority of gene expression variation is not due to genomic changes but is attributable to differences between patients or other unaccounted variables.

Indeed, when we used GSEA NES as a metric for detection sensitivity (see Fig. 7), DEGs associated with PTEN loss benefitted most from an LMM analysis that was capable modeling the intra-patient or intra-lesion variability as a blocking effect. In addition, the co-occurrence of relatively homogeneous *TMPRSS2-ERG* fusion and PTEN loss within each focus from cohort 1 created additional challenges during the *PTEN* loss analysis for the four aggregation methods but were significantly reduced by LMM. Thus, inclusion of the foci variability using models such as LMM significantly increased sensitivity and specificity of an analysis to arrive at per-patient phenotypes. While the addition of *TMPRSS2-ERG* status as a parameter in a *PTEN* loss analysis could mitigate the confounding effect of *TMPRSS2-ERG* fusion, doing so in standard practice would require its status known *a priori*, and patient level transcriptomic studies are often complex and thus the inclusion of such parameter would not be always possible.

Compared to cohort 1, which was comprised of laser capture microdissected tumor foci, cohort 2 was intrinsically more heterogeneous within-foci. The approach used for sampling prostate tumors used in cohort 2 more closely mirrors that used in clinical settings, whereby a large region expected to represent the most aggressive region is generally selected for molecular profiling [28]. Thus, when more than one area was selected in this manner, the variance in DEG profiling the entire patient was distinctive with an LMM, whereas averaging/summing counts or FASTQ concatenation arrived at similar profiles. However, when we consider NES as ground truth for this comparison, generalized to all three target genotypes no single method outperformed the other. Nonetheless, the advantage to LMM is the use of blocking factors for fixed effects and the ability to assess within-lesion or within-patient heterogeneity against one or more salient phenotypes.

## CONCLUSIONS

The aggregation of multiple transcriptomes from a single tumor to estimate per-patient phenotypes will vary in its reliability as a direct function of heterogeneity. Tumors often harbor heterogeneous alterations within the same geographic region, resulting in vastly different gene expression profiles between nearly-adjacent tumor cells. Experimental designs that include tumor resampling have improved power to detect driver genotypes and the variance of gene expression or biomarker profiles related to phenotype. Although we found that a linear mixed model was a robust aggregation method that performed consistently well in analyses with homogeneous and heterogeneous phenotypes within individual tumors, patient-level summarization methods including RNA-seq count averaging and summation showed benefit for detecting gene expression changes that were associated with more homogeneous genotypes. Consequently, the method of per-patient gene expression summarization is highly context dependent on any salient target genotypes and the tolerance of variability for formulating robust conclusions.

## KEY POINTS

- Within- and between-lesion tumor heterogeneity can exist within the same patient. Strategies to sampling the same tumor multiple times can improve detection of intratumoral phenotypic variability.
- The performance of different methods to aggregate multiple transcriptomes into per-patient summaries was highly dependent upon the phenotype of interest.
- Dominant or clonal transcriptomic signals in prostate cancer, such as the *TMPRSS2-ERG* fusion, were robust to multiple aggregation methods, although subclonal alterations such as *PTEN* loss were more conducive to linear mixed models.
- Comparison of different aggregation methods to benchmarked gene expression profiles can identify unappreciated sources of within-patient transcriptomic heterogeneity based on defined genomic alterations.

## DATA AVAILABILITY

The data underlying this article are available in:

- dbGaP at https://www.ncbi.nlm.nih.gov/gap/, and can be accessed with accession ID phs001938.v2.p1; and
- SRA at https://www.ncbi.nlm.nih.gov/sra, and can be accessed with accession ID PRJNA579899.
- NCI GDC at http://portal.gdc.cancer.com, and can be accessed with the project ID TCGA-PRAD.

## ACKNOWLEDGMENTS

Portions of this work utilized the computational resources of the NIH HPC Biowulf cluster.

## SUPPLEMENTARY INFORMATION

Supplementary Table 1: DEGs for *TMPRSS2-ERG* fusion from cohort 1

Supplementary Table 2: DEGs for *PTEN* loss from cohort 1

Supplementary Table 3: DEGs for 18q loss from cohort 1

Supplementary Table 4: DEGs for *TMPRSS2-ERG* fusion from cohort 2

Supplementary Table 5: DEGs for *PTEN* loss from cohort 2

Supplementary Table 6: DEGs for 18q loss from cohort 2

Supplementary Table 7: The top 75 up-regulated genes for each gene set

## Notes

**COMPETING INTERESTS** The authors declare no competing interests.

**FUNDING** This work was supported by the Prostate Cancer Foundation (Young Investigator Awards to S.W. and A.G.S.), the Department of Defense Prostate Cancer Research Program (W81XWH-19-1-0712 to S.W., W81XWH-16-1-0433 to A.G.S.), and the Intramural Research Program of the NIH, National Cancer Institute.

https://www.ncbi.nlm.nih.gov/gap/

https://www.ncbi.nlm.nih.gov/sra

### Competing Interest Statement

The authors have declared no competing interest.

